# R2DGC: Threshold-free peak alignment and identification for 2D gas chromatography mass spectrometry in R

**DOI:** 10.1101/179168

**Authors:** Ryne C. Ramaker, Emily Gordon, Sara J. Cooper

## Abstract

**Summary:** Comprehensive two dimensional gas chromatography-mass spectrometry is a powerful method for analyzing complex mixtures of volatile compounds. This method produces a large amount of raw data that requires downstream processing to align signals of interest (peaks) across multiple samples and match peak characteristics to reference standard libraries prior to downstream statistical analysis. To address the paucity of applications addressing this need, we have developed an R package that implements retention time and mass spectra similarity threshold-free alignments, seamlessly integrates retention time standards for universally reproducible alignments, performs common ion filtering, and provides compatibility with multiple peak quantification methods. We demonstrate the packages utility on a controlled mix of metabolite standards separated under variable chromatography conditions and data generated from cell lines.

**Availability and documentation:** R2DGC can be downloaded at https://github.com/rramaker/R2DGC or installed via the Comprehensive R Archive Network (CRAN).

**Supplementary information:** Supplementary data are available at Bioinformatics online.

## 1 Introduction

Metabolomics is a rapidly growing field that seeks to comprehensively measure a set of small molecule metabolites in a sample (Wishart, 2016). Two dimensional gas chromatography-mass spectrometry (2D-GCMS) was developed to improve chromatographic separation by allowing for improved measurement of complex mixtures from biological samples (Mondello *et al.*, 2008). 2D-GCMS couples two gas chromatography (GC) columns with complementary chemistries, to a time-of-flight mass spectrometer. The resulting data contain three identifying components for each feature (metabolite): two retention times, one from each chromatographic separation, which reflect the affinity of a compound to each column, and a mass spectrum that is relatively unique to each compound. Each component has associated error. Retention times can vary across experiments or instruments due to slight differences in chromatography methods or slight variations in column length, chemistry, and age. Furthermore, subjecting metabolites to electron spray ionization, the most common ionization method for 2D-GCMS, can lead to similar mass spectras among structurally related compounds. However, combining these three components provides a powerful method for deconvoluting complex mixtures.

Metabolomics data processing and analysis can be divided into three steps. First, raw signal is processed to distinguish analyte peaks from background noise and relative peak areas and spectra are computed. The next step involves alignment of peaks common to a group of samples and identification of metabolites from which peaks were likely derived by matching to a reference library. Lastly, statistical methods are used to compare peak areas or metabolite concentrations between groups of samples. Here, we describe a novel software package to streamline the second data processing step of peak alignment and metabolite identification. We also provide a retention indexed reference library containing information on 298 peaks derived from over 125 metabolite standards and commonly observed background peaks for use with the package.

## 2 Methods

R2DGC is designed to take individual sample files containing basic peak information and perform pre-processing steps, generate an alignment table with areas of peaks common to multiple samples, and match aligned peaks to a reference library (Fig. 1). Detail on algorithms used by the package is provided in the Supplemental Material.

**Fig. 1.**
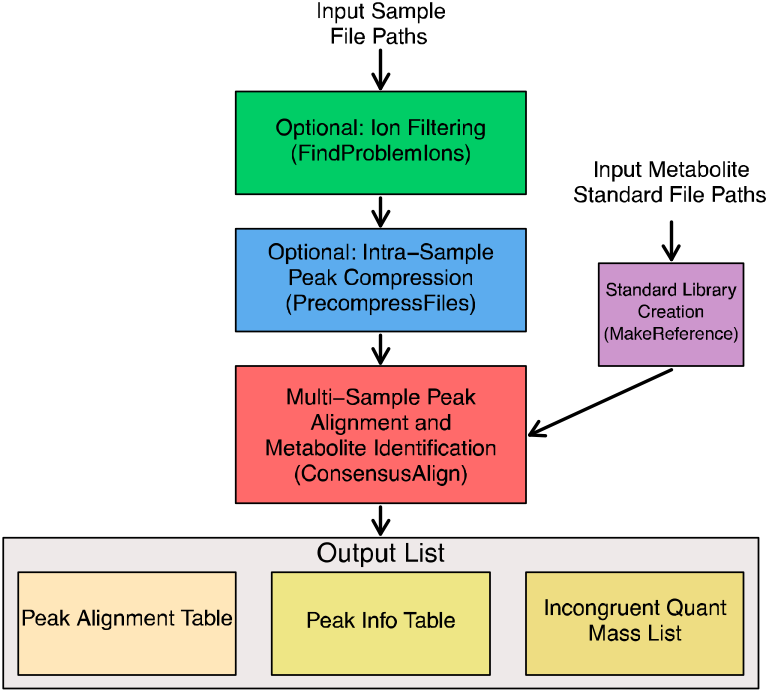
R2DGC alignment pipeline workflow

### Input data

The input for peak alignment with R2DGC is a list of file paths to individual sample files that contain the retention times, area, and mass spectra of each peak as commonly outputted by vendor software like ChromaTOF (Supplemental Section 2.2). The user can also specify the peak quantification method used to ensure proper downstream processing. For example, in order to ensure accurate quantification, the aligner will ensure all aligned peaks peak areas were computed with the same quant mass. If quant mass incongruencies exist, the user is notified and a preliminary peak area conversion is performed. If primary or secondary retention time standards were used, the user simply needs to ensure a consistent naming scheme is used to identify each standard peak and make use of retention time indexing.

### Filtering common ions

The first optional pre-processing function (FindProblemIons) allows users to filter ions from the mass spectra of each peak to improve downstream spectral matching accuracy and efficiency. Ions that are found to be absent from all peaks or are products of the derivatization protocol common to all peaks are identified for removal (Supplemental Section 2.3). This function, as well as those described below, takes advantage of parallel computing in R to ensure alignments efficiently scale with increasing sample sizes.

### Intra-sample peak compression

The second optional pre-processing function (Precompress Files) searches each sample file for peaks that were incorrectly split in primary peak identification and quantification and likely represent the same analyte. This often occurs due to signal saturation for abundant metabolites and can cause problems in downstream alignments because peaks will be assigned to only one of the split peaks leaving apparently missing data in the final alignment table (Supplemental Section 2.4). Manually identifying these peaks in each sample is an extremely time intensive process.

### Generating a standard library

The third pre-processing function (MakeReference) allows users to generate a reference standard library to identify metabolites from which aligned peaks are likely derived. The user supplies a list of file paths to metabolite standard files containing peak retention time and mass spectra information. If retention time standard indexing is desired, the user also includes retention time standard peaks with each metabolite standard, using a naming scheme identical to the sample file input described above (Supplemental Section 2.5). We have provided a pre-formatted standard reference library containing greater than 125 metabolites each with their primary retention time indexed with nine fatty acid methyl esters (FAMEs) that is installed with the package.

### Multi-sample peak alignment

The final and only required function (ConsensusAlign) takes sample files and an optional metabolite standard library and aligns common peaks across samples. It then identifies likely metabolites from which each aligned peak was derived. The aligner computes pairwise similarity scores between each peak in each sample file to each peak in a seed file. Similarity scores are computed as the dot product of each mass spectra penalized by differences in peak retention times. An optimal similarity score threshold (maximizing the number of peaks with at least one match while minimizing the total number of potential matches) for peak alignment is computed prior to aligning peaks. Alternatively, the user can manually alter the stringency of the alignment. After primary alignment, an optional second alignment can be performed that searches for ‘high likelihood’ missing peaks that are present in nearly all samples at a slightly relaxed similarity threshold. To ensure the seed file selected does not bias the alignment, the user can elect to use multiple seed files and retain only peaks aligned by a majority of seed files. This function outputs an alignment table with peak areas for each sample (columns) by each aligned peak (row), a peak information table with retention times, mass spectra, reference library matches for each peak, and, depending on the peak quantification method used, a list of peaks that were aligned, but whose peak areas were computed using different masses (Supplemental Section 2.6).

## 3 Results

We have extensively tested the package on both known mixtures of metabolite standards, and extracts derived from tissues and cell lines. We provide results from two such test cases to demonstrate the utility of the package (https://github.com/rramaker/R2DGC_Datasets). The first test case involves six total samples representing three different known mixtures of amino acids analyzed with two different chromatography methods (Supplemental Section 4.1). The second is a more typical metabolomics test set that includes 24 samples from pancreatic cancer cell lines for which 12 amino acids of each sample were manually aligned to benchmark aligner performance (Supplemental 4.2). We find our alignment pipeline performs comparably to two previously developed open source alignment pipeline (Castillo *et al.*, 2011; Jeong *et al.*, 2012) often resulting in more true alignments and fewer incorrect or missing alignments.

## 4 Conclusions

R2DGC provides a comprehensive, efficient, and reproducible pipeline for 2D-GCMS peak alignment and identification. It is one of the few freely available and actively maintained 2D-GCMS aligners currently available and improves upon existing software by (1) calculating optimal retention time or mass spectra similarity thresholds rather than an arbitrary designation, (2) facilitating the use of retention time standards to allow for universally reproducible alignments, (3) filtering ions that hinder mass spectra comparisons, (4) and providing compatibility for multiple peak quantification methods.

